# Thermally Reversible Pattern Formation in Arrays of Molecular Rotors

**DOI:** 10.1101/2022.10.19.512893

**Authors:** Marcello DeLuca, Wolfgang G. Pfeifer, Benjamin Randoing, Chao-Min Huang, Michael G. Poirier, Carlos E. Castro, Gaurav Arya

## Abstract

Control over the mesoscale to microscale patterning of materials is of great interest to the soft matter community. Inspired by DNA origami rotors, we introduce a 2D nearest-neighbor lattice of spinning rotors that exhibit discrete orientational states and interactions with their neighbors. Monte Carlo simulations of rotor lattices reveal that they exhibit a variety of interesting ordering behaviors and morphologies that can be modulated through rotor design parameters. The rotor arrays exhibit diverse patterns including closed loops, radiating loops, and bricklayer structures in their ordered states. They exhibit specific heat peaks at very low temperatures for small system sizes, and some systems exhibit multiple order-disorder transitions depending on inter-rotor interaction design. We devise an energy-based order parameter and show *via* umbrella sampling and histogram reweighting that this order parameter captures well the order-disorder transitions occurring in these systems. We fabricate real DNA origami rotors which themselves can order *via* programmable DNA base-pairing interactions and demonstrate both ordered and disordered phases, illustrating how rotor lattices may be realized experimentally and used for responsive organization. This work establishes the feasibility of realizing structural nanomaterials that exhibit locally mediated microscale patterns which could have applications in sensing and precision surface patterning.

## 1. INTRODUCTION

The Ising model^1^ has long served as a valuable pedagogical device for describing order-disorder transitions. This model, originally developed to explain ferromagnetism, treats magnetic spins as one of two states: spin up or spin down, where it is energetically favorable for neighboring spins to be pointing in the same direction. In a one-dimensional lattice of spins, the enthalpic cost of disorder becomes negligible in the thermodynamic limit *N* → ∞, so the system will be disordered at any temperature *T* > 0. However, in dimensions two or higher this enthalpic penalty to disorder becomes non-negligible, which gives rise to a first-order phase transition between an ordered and a disordered state at finite temperatures. This model enables many features of phase transitions to be captured, including spinodal and binodal decomposition as well as divergent magnetic susceptibility and correlation length. Furthermore, other transition types such as gas-liquid transitions^2^ and mixing-demixing transitions^3^ are also directly mappable to the Ising model, making it useful for studying real physical systems in addition to hypothetical ones.

Order-disorder transitions may be broadly classified into two categories: those that involve translational motion of the system constituents, either between phases of different densities (e.g., in gas-liquid transitions) or local compositions (e.g., in mixing-demixing transitions), and those that involve only local rotational order and whose phase transitions incur no change in density or composition (e.g., magnetic spins). Translation-based phase transitions are ubiquitous both in nature and in engineering systems, where they have been harnessed for numerous applications. For instance, in metallurgy, specific thermal annealing procedures can yield distinct phases of steel which have drastically different mechanical properties, a phenomenon that has been exploited for thousands of years.^4^ However, to the best of our knowledge, orientation-based constant density phase transitions have not been exploited in rationally designed nanomaterial systems.

DNA origami^5^ provides an opportunity to realize such a system using local interactions between neighboring structures to induce large-scale control over ordering as is seen in magnetic systems. DNA origami is a technique for rationally designing nanostructures using DNA as a building material. This allows DNA to be used as a building block for constructing arbitrary, yet precise, shapes and patterns. Many dynamic devices have been made from DNA,^6-8^ including rotary devices,^9-12^ which allows one to imagine the construction of an array of nanomechanical rotors which could exhibit neighbor interactions which may drive constant density phase transitions fundamentally similar to those in conventional spin lattices. Importantly, at the length scales of DNA origami, thermal fluctuations still play a key role in organizational behavior, and order-disorder transitions may still be directly observable using standard imaging techniques such as transmission electron microscopy (TEM) or atomic force microscopy (AFM).

Here, we introduce a new class of self-organizing lattice systems that we call “rotor lattices” (distinct from another lattice model of the same name^13^) which exhibits unique behavior such as low transition temperatures and multiple order-disorder transitions and can be realized as physical 2D arrays of DNA origami rotors using base pairing interactions between complementary single-stranded oligonucleotides (sticky ends)^14-16^ or unstacked bases at the ends of DNA helices (blunt ends)^17-21^ to mediate their organization. These rotor lattices are motivated by phase transitions of water where the geometric nature of neighbor interactions between molecules dictates their transition behavior and ordering in liquid and solid phases.^22^ 2D lattices of rotors should be simple to image and provide a 2D model system which could eventually inform the organization of 3D systems. Rotary elements can only form a finite number of bonds, which can give rise to behaviors such as termination and the formation of switchable shapes and patterns that can be toggled between their phases by modulating the interaction strength. We use Monte Carlo simulations to predict the unique organizational behavior of these rotor lattices, and then demonstrate how DNA origami may be used to actualize such a system by designing small arrays of DNA rotors which exhibit nearest-neighbor interactions through overhang base pairing. The DNA rotors are shown to exhibit both ordered and disordered phases, establishing the feasibility of realizing Ising-like structural nanomaterials.

## 2. RESULTS

### Model Development

The magnetic lattice models that inspired this work normally fit within the *n*-vector model^23^ wherein a *d*-dimensional lattice of spins, which have their own dimensionality *n*, interact with all of their nearest neighbors. In the case of the Ising model, *n* = 1 and each site contains a 1D spin. The *n* = 2 case describes the XY model,^24^ and the *n* = 3 case, known as the Heisenberg model,^25^ is used to study phenomena such as ferromagnetism. The rotor lattice model that we describe here does not fit within the *n*-vector model. It instead utilizes directional interactions where an interacting “arm”, which protrudes from the rotor, can interact with only a *single* neighbor, contrasted by the spins’ ability to interact with all neighbors in the *n*-vector model or in variations of the Ising model such as the Potts model.^26^ We will show that this gives rise to its own class of interesting transition behaviors.

Our rotor lattice model is described by a space-filling unit cell in any number of dimensions *d* and coordination number *z*. Each unit cell harbors a single rotor which can exhibit *p* orientations coinciding with the directions of neighboring unit cells (in the case of 2D systems, *p* can equal some natural number multiple of *z*, but in 3D systems, this becomes more complex, e.g., a 3D cubic rotor in a 3D cubic unit cell can exhibit 24 or more orientations). Each “face” of the rotor contains an interacting region (arm) that can interact with a neighboring rotor in the direction the arm is pointing. In this work, we will focus on the *d* = 2 and *p* = *z* = 4 case, i.e., a 2D square lattice with four-armed rotors (**Fig. 1a**). The rotors can be diagrammatically described using squares with colored arms sticking out in each interaction direction (**Fig. 1b**). Same-colored arms that point toward each other interact with an energy corresponding to the lesser of the widths of the two interacting arms (**Fig. 1c**), so, a rotor can only exhibit favorable interactions with another rotor if both possess arms of the same color and non-zero width along their coordination direction. For example, a rotor face with one sticky end can interact with a rotor face with three identical sticky ends, but only one of those three sticky ends can hybridize and thus the energy change induced by this interaction will only correspond to base pairing between a single set of sticky ends. Such interaction scenarios could be realized experimentally in DNA origami using blunt ends or sticky ends of varying strength or quantity.

**Figure 1:**
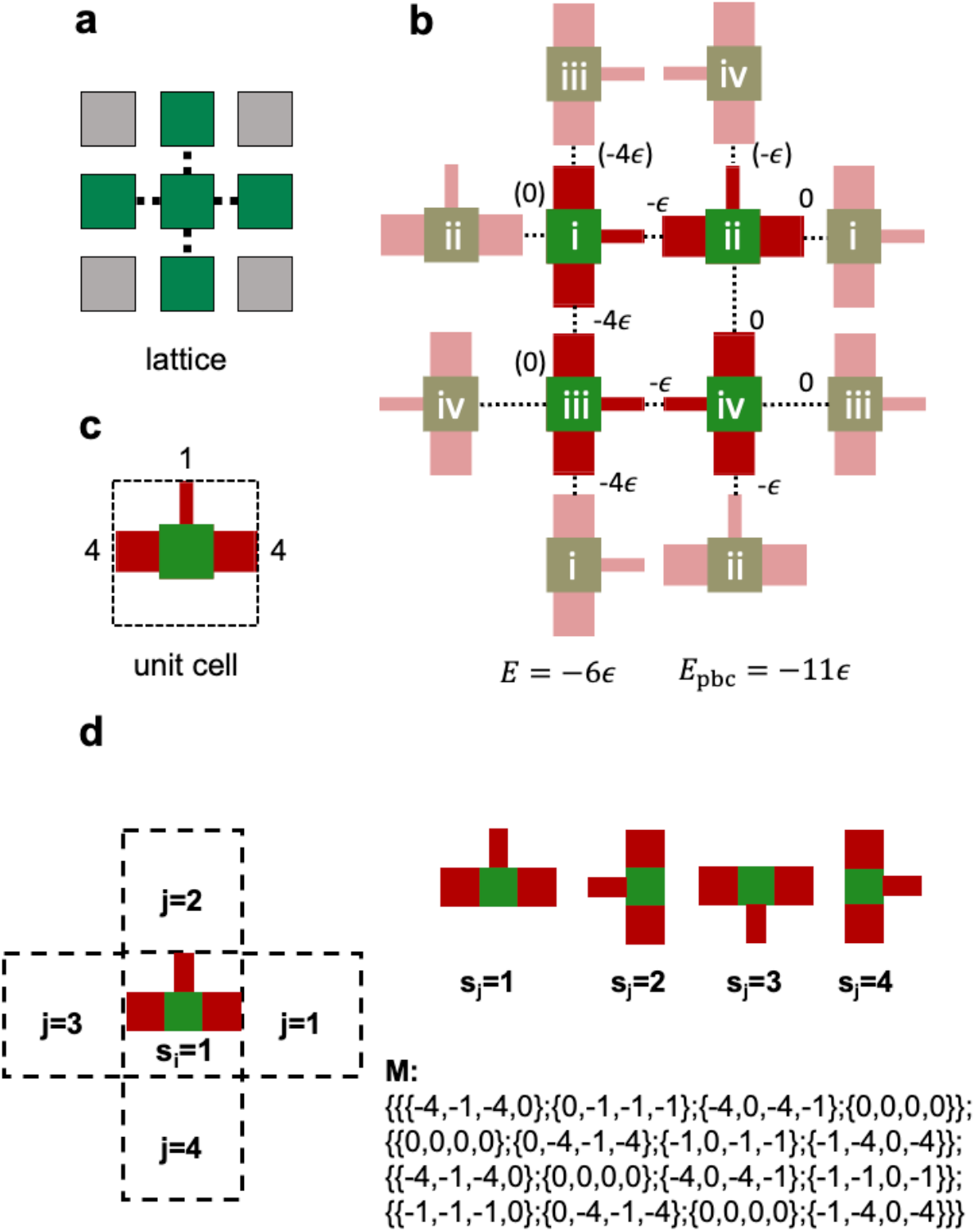
Rotor lattice model illustrating the lattice structure with square interactions (a), the rotor element in each unit cell (b), energy evaluation for an example 2×2 system with and without periodic boundaries (c), and the interaction tensor used for describing every possible interaction (d). Light-colored rotors represent mirror images from the minimum image convention. Energies in parentheses have already been accounted for on the opposite boundary and are not added to the microstate energy.

Quantifying the energy of microstates of such a rotor lattice system requires the use of a Hamiltonian containing a 3D interaction tensor **M** of dimensions *p* × *z* × *p* which accounts for the orientational state of the rotor, the coordination site of the rotor’s neighbor, and the orientational state of the neighbor:

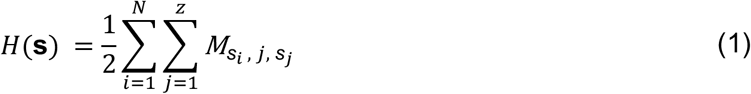

where **s** is the system microstate containing the states of *all* rotors, *N* is the number of rotors in the device, *z* is the number of neighbors that each device has (coordination number), s_*i*_ is the orientational state of one device, *j* is the site location of the neighbor being compared, and s_*j*_ is the orientational state of that neighbor. In **Fig. 1d**, we provide an example of an interaction tensor corresponding to the rotor design shown in the figure, where all interaction strengths are specified in units of *ϵ*, the energy scale of the system. Interaction tensors for all other rotors in this study can be found in the SI in **Fig. S1-S5**.

Standard Monte Carlo (MC) simulations can be used to investigate the ordering behavior of rotor lattices at equilibrium for specified reduced (dimensionless) temperature *k*_B_*T*/*ϵ*, which could be tuned in the real system by varying the actual temperature or the environmental conditions such as salt concentration or pH that modulate *ϵ*. We employed a single MC move type consisting of random rotations to one of the other *p* − 1 orientations of randomly selected rotors (from *N* rotors) with the standard Metropolis acceptance criterion^27^:

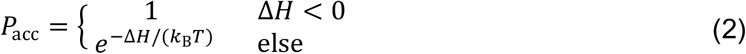

where Δ*H* is the change in the lattice energy described by Eq. 1 and *k*_B_ is the Boltzmann constant. We found that this single move type was sufficient to adequately sample the microstates of the rotor lattice for the range of temperatures used in this work.

Using this simulation scheme, we studied the ordering behavior of six distinct rotor designs, including two that exhibit Ising-like behavior in 1D and 2D (**Fig. 2** and **Fig. S6**):

**Figure 2:**
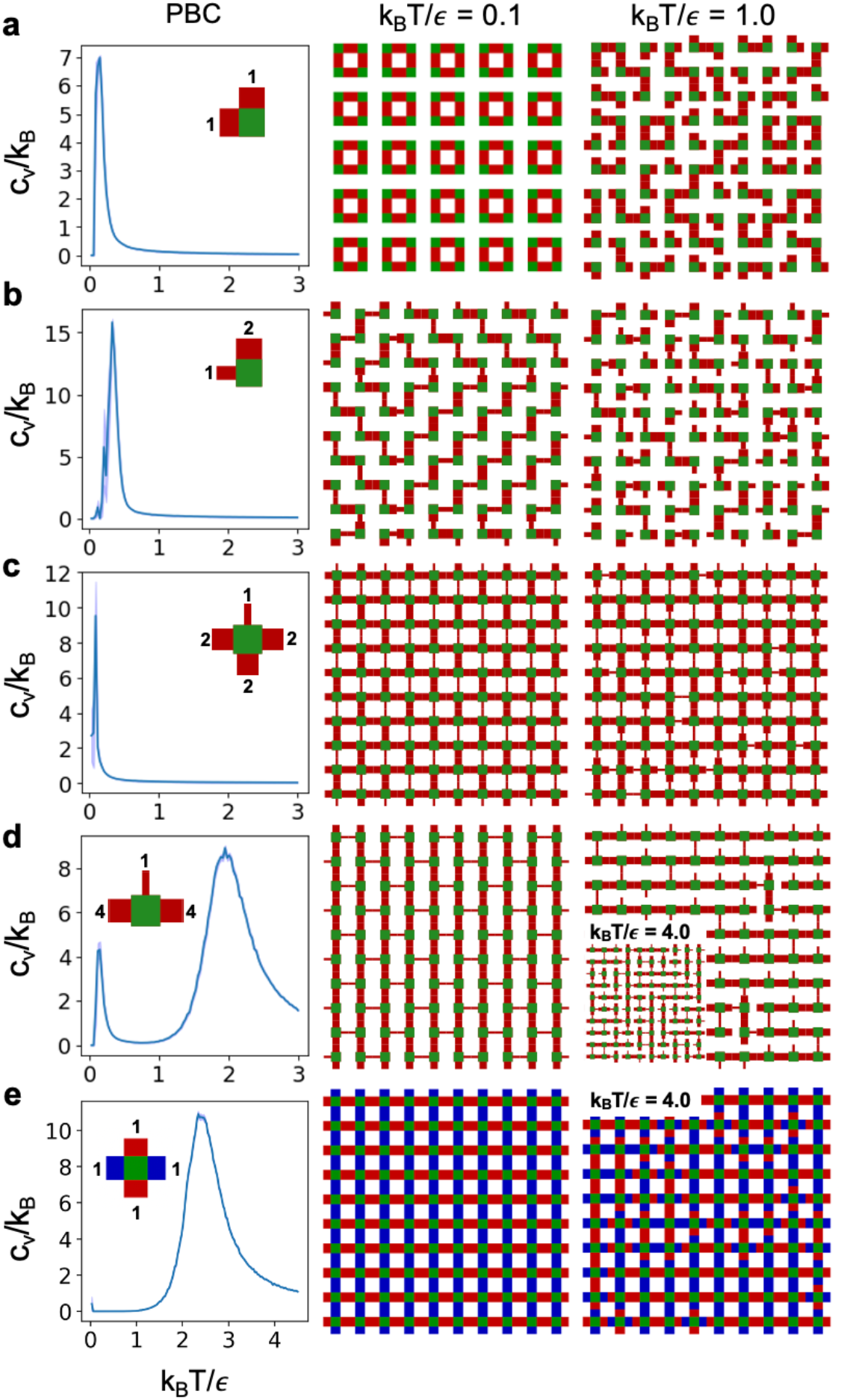
Specific heat behavior of non-canonical spin systems and some representative examples of array microstates: (a) L-shaped rotor. (b) Chiral rotor. (c) 2-2-2-1 rotor. (d) 4-1-4 rotor. (e) Ising-like rotor.

- *L-shaped rotor* contains two arms each of interaction strength −*ϵ* located at a 90° angle to each other
- *Chiral rotor* has the same arm locations as the L-shaped rotor but with one arm’s strength double relative to the other, that is, arms of strengths −*ϵ* and −2*ϵ*
- *2-2-2-1 rotor* containing three stronger arms of strength −2*ϵ* and a single weak arm of relative strength −*ϵ*
- *4-1-4 rotor* contains two opposite stronger arms of relative strength −4*ϵ* and a single weak arm of relative strength −*ϵ*
- *2D Ising-like rotor* contains two different types of arms that interact favorably with arms of their type (with energy −*ϵ*) and unfavorably with arms of the other type (with energy +*ϵ*)
- *Quasi-1D Ising rotor* contains two arms on opposite faces of interaction strength −*ϵ*

In each case, we examined 2D square lattices of rotors of size 10×10, with and without periodic boundary conditions (PBCs).

### Order-Disorder Transitions of Rotor Lattices

Phase transitions are usually characterized by a peak in specific heat at some temperature. The reason this occurs is that the average microstate energy of the disordered phase is significantly higher than the average microstate energy of the ordered one; the specific heat peak arises because the system is able to transition back and forth between relatively ordered and relatively disordered states as the energy barrier between two states disappears at this transition (critical) temperature, denoted by *T*_c_. Therefore, we carried out MC simulations for each of the six rotor designs introduced earlier and the specific heat was determined from the energy fluctuations:

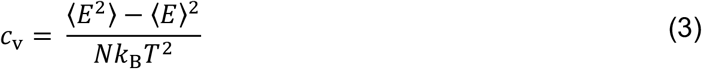

where ⟨·⟩ denotes an ensemble average. By plotting *c*_v_ as a function of *T*, we can then use local maxima in specific heat to identify order-disorder transitions.

**Figure 2** shows the heat capacity curves for several of the studied rotor systems. All systems show a peak in the specific heat when using periodic boundary conditions, indicating that they all exhibit order-disorder transitions, though the number of such transitions and the transition temperature varies across the six rotor designs. Specifically, the L-shaped, chiral, 2-2-2-1, and quasi-1D rotors exhibit peaks at low temperatures (*k*_B_*T*/*ϵ* < 1), significantly lower than those expected from first-order phase transitions associated with the Ising lattice (**Fig. 2a-c and Fig. S7**); the 2-2-2-1 rotor with very small differences between all interactions exhibits an especially low transition temperature of *k*_B_*T*/*ϵ* ≈ 0.1. The 2D Ising-like rotor (**Fig. 2e**), as expected, exhibits a peak at the same temperature as the classic 2D Ising lattice (*k*_B_*T*/*ϵ* ≈ 2.269). Lastly, and most intriguingly, the 4-1-4 rotor exhibits two peaks, one at a very low temperature, akin to those of the L-shaped and chiral rotors, and another at a much higher temperature which is coincidentally closer to the *T*_c_ of the classic 2D Ising lattice (**Fig. 2e**).

All non-Ising rotor designs form interesting morphologies at low temperatures in the ordered phase. For example, the L-shaped rotor tends to form arrays of closed loops (**Fig. 2a**). Increasing the strength of interactions on one arm to form the chiral rotor leads to the formation of chiral zigzags or radiating loops (**Fig. 2b**). The 2-2-2-1 rotor forms a brick wall structure from the three stronger arms’ interactions (**Fig. 2c**). The 4-1-4 rotor (**Fig. 2d**) tends to form an alternating ladder-like morphology at very low temperatures, while at intermediate temperatures, the “rungs” of this morphology will tend to dissociate while the stronger interactions will remain intact. At the highest temperatures, all order is lost. Such systems thus exhibit three distinct morphologies; correspondingly, this design exhibits multiple peaks in the specific heat curve.

The critical temperature of each order-disorder transition can be qualitatively explained by the enthalpic difference between ordered and disordered phases. For each design, the ordered energy ⟨*E*_*T*→0_⟩ is equal to that of the ground state, which may be directly enumerated or can be computed using MC simulations. With PBCs, the disordered energy reflects a completely random selection of microstates and can be shown to reduce to:

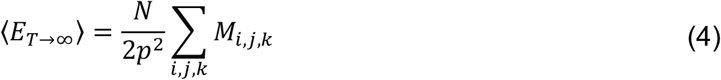

where ⟨*E*_*T*→0_⟩ is the average microstate energy as temperature approaches infinity. The Ising lattice has an energy difference between ordered and disordered states of −2*ϵ* per unit and a transition temperature *k*_B_*T*/*ϵ* ≈ 2.269. The L-shaped rotor has an energy difference between ordered and disordered states of −0.5*ϵ* per unit and a transition temperature that is correspondingly much smaller, *k*_B_*T*/*ϵ* ≈ 0.25. The 2-2-2-1 rotor has an even smaller energy difference between ordered and disordered states of −0.375*ϵ* and a correspondingly even smaller transition temperature *k*_B_*T*/*ϵ* ≈ 0.1. Assuming that the intermediate phase of the 4-1-4 rotor retains strong bond order but destroys weak bond order, this system has an energy difference between the lowest temperature and intermediate temperature phases of −0.5*ϵ* per unit and correspondingly has a similar first transition temperature to the L-shaped system, while the difference between intermediate and disordered phases is −1.375*ϵ*, yielding a much higher transition temperature than the other systems, except for the Ising lattice. It is noted that the lack of perfectly linear scaling between transition temperature and the energy difference between ordered and disordered states likely arises from multiple factors including entropy and the cooperative nature of these geometric bonds, i.e., the ability of bonds to form connected chains.

Also of interest in these systems is the dependence of the height of the specific heat peaks on the presence or absence of PBCs. While most systems tend to form structures whose interactions span the entire array and therefore have higher specific heat peaks when PBCs are used (see **Fig. S7**), the L-shaped rotor array exhibits a lower peak with PBCs. This rotor prefers to form closed loops in its ordered state, so the more disordered state is in fact easier to achieve with periodic boundaries because the entropy increase of the disordered phase in this system is significant when PBCs are used and there is no enthalpic benefit to PBCs in any system with even-sized arrays, while the enthalpic benefit of PBCs outweighs the entropic penalty of order for all other systems at low temperatures.

### Order Parameters

In contrast to magnetic systems which can easily be described by average magnetization, there is no simple analogous quantity for characterizing order in spins with four orientational states. Since the microstate energy is a function of system configuration and number of bonds, we elect to use a microstate energy-based order parameter to describe order-disorder transitions:

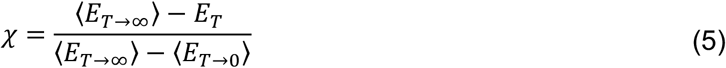

This equation ensures that the order parameter is 1 at zero temperature where the system resides in the lowest energy microstate, and the order parameter is 0 as temperature approaches infinity where there is effectively no bias toward any microstate. **Figures S8 and S9** demonstrate that this order parameter describes order-disorder transitions as well as the magnetization parameter in the case of the Ising lattice, suggesting that an energy-based order parameter may also be sufficient for describing order-disorder transitions in other rotor systems.

To obtain the instantaneous order parameter *χ* for any microstate using Eq. 5, we need to first obtain the two limits ⟨*E*_*T*→0_⟩ and ⟨*E*_*T*→∞_⟩. At low temperatures, most systems have a trivial ground state and ⟨*E*_*T*→0_⟩ can be readily enumerated from this state. Where systems do not have a trivial ground state, ⟨*E*_*T*→0_⟩ can be computed using MC simulations. At high temperatures, ⟨*E*_*T*→∞_⟩ may be analytically obtained using Eq. 4. To characterize the ordering behavior of rotor lattices as a function of temperature, we conducted umbrella sampling^28^ MC simulations on four different 10×10 rotor designs using *χ* as a reaction coordinate. Using weighted histogram analysis, we then determined the free energy of the system as a function of *χ*. The value of *χ* corresponding to the lowest free energy at any temperature, *χ*_min_, provides the most representative value of order at a given temperature.

**Figure 3** shows these obtained values of *χ*_min_ as function of reduced temperature *k*_B_*T*/*ϵ* for four of our rotor designs as a way of characterizing their ordering behavior. We found that the L-shaped, quasi-1D Ising, and Ising-like systems exhibit a single inflection point approximately coinciding with the temperature of their specific heat peak (**Fig. 3b-d**), while the 4-1-4 design exhibits three inflection points corresponding to its two peaks in specific heat (**Fig. 3a**). These results indicate that this order parameter is reasonably capturing the order-disorder transition behavior of this system. The more gradual slope of the 4-1-4 rotor design is also consistent with its broader specific heat peak compared to the other rotor designs.

**Figure 3:**
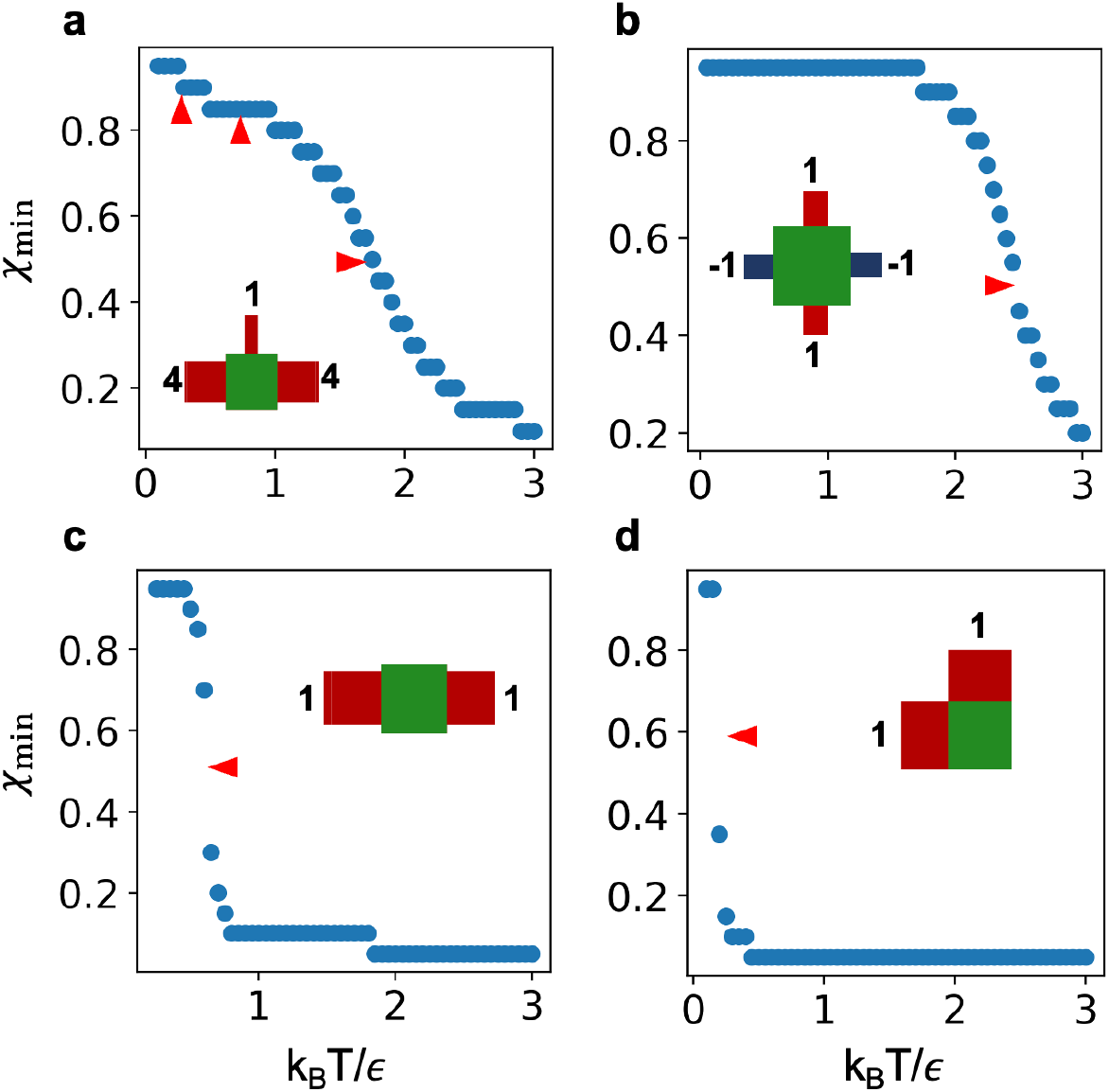
Free energy minimizing order parameters as a function of reduced temperature for four different rotor designs: (a) 4-1-4 rotor, (b) Ising rotor, (c) quasi-1D Ising rotor, and (d) L-shaped rotor. All sampling was performed for 10×10 lattices using PBCs. Approximate locations of the inflection points are shown with red arrowheads.

To investigate the role of specific arms in order-disorder transitions, especially those with multiple order-disorder transitions, we can additionally assign a single order parameter *ψ* to each arm of a rotor design:

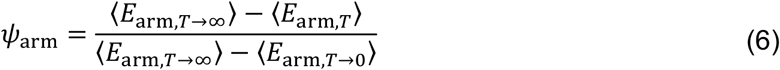

where *E*_arm_ is the binding energy associated with a particular arm of the unit cell. This discretizes the order parameter into distinct components that help identify anisotropy and the specific sites involved in order-disorder transitions.

In the case of the Ising-like system, all four arms lose order simultaneously (**Fig. 4d**); the same is true of the L-shaped and quasi-1D Ising rotor designs (**Fig. 4b-c**). However, the “4-1-4” design loses order in the weak arm first and then the stronger arms at a higher temperature (**Fig. 4a**). Also interesting about this design is that the weak arms are less ordered in the intermediate state than in the fully disordered state, yielding a negative order parameter. This is due to the persistence of the stronger bonds leaving the smaller ones in unfavorable configurations, while at higher temperatures, the small arm has the opportunity to bond with any of the three rotor arms, yielding an order parameter approaching zero. This corroborates our observation of three distinct phases for this system and emphasizes the role of individual arm order in its transitions.

**Figure 4:**
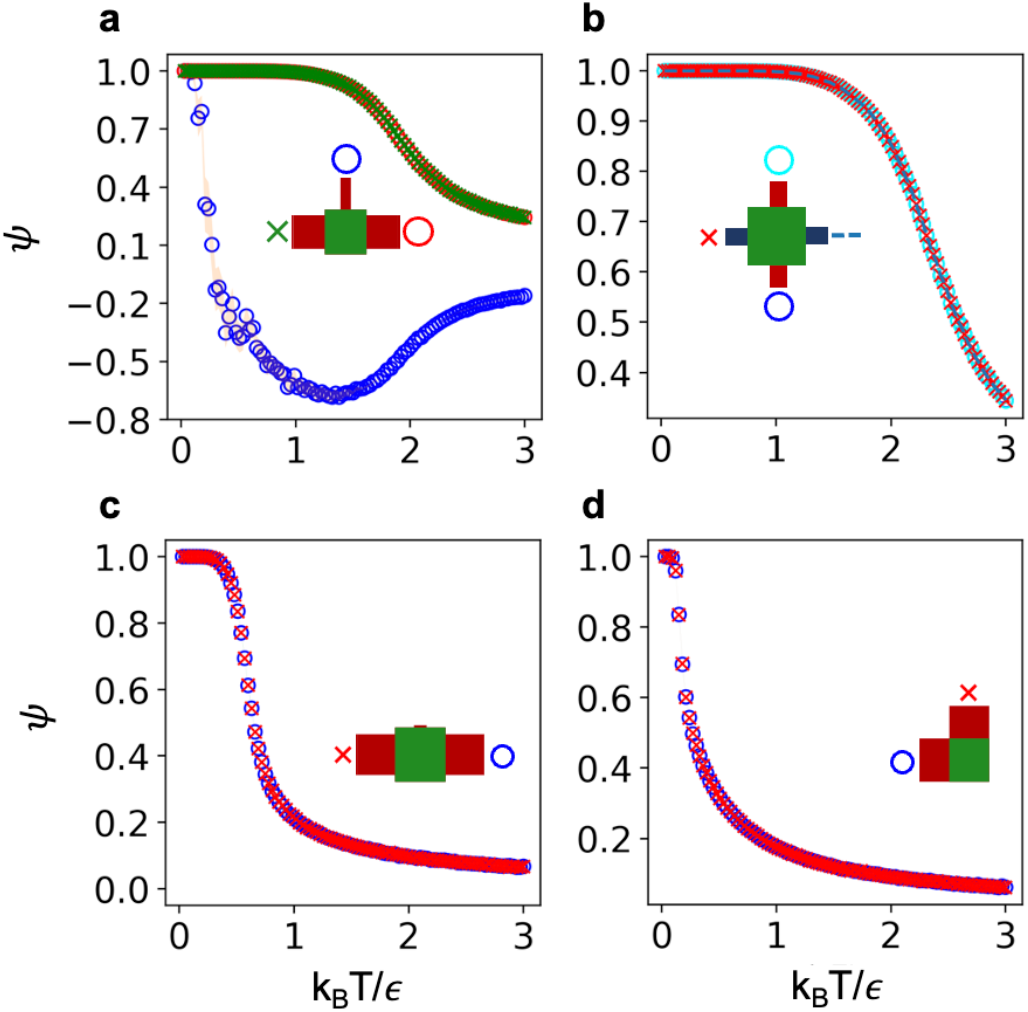
Arm-specific order parameters as a function of reduced temperature for: (a) 4-1-4 rotor, (b) Ising rotor, (c) quasi-1D Ising rotor, and (d) L-shaped rotor.

### Experimental design of rotor arrays

To demonstrate the feasibility of experimentally realizing rotor arrays, we designed and fabricated DNA origami nanostructure (DON) designs with rotary elements coupled to a fixed base platform capable of self-assembling into arrays. These structures were designed using the MagicDNA design software package.^29^ DNA origami structural units were designed to contain two L-shaped rotor elements each to enable a simpler assembly of 2×2 arrays. We used hierarchical assembly protocols^30^ to form the two-rotor unit into 2×2 L-shaped rotor arrays. The two-rotor structural units were folded from two orthogonal sequence scaffolds in a single-pot folding reaction. We first tested folding over a broad range of magnesium concentrations (**Fig. S10**). Based on this initial screen, we chose to use 14 mM MgCl_2_ to fold the two-rotor units. TEM imaging of gel-purified structures revealed that they were well-folded and properly attached to the base platform, containing two randomly oriented rotors in the absence of specific inter-rotor binding interactions (**Fig**.**5b, Fig. S11**). To form the 2×2 array, we folded two separate versions of the two-rotor unit, each version being passivated on one side, either left or right, of the base platform with poly-T overhangs and with unpaired scaffold strand on the other side, allowing for selective higher order assembly only on one side leading to dimer formation. After removal of excess staple strands, dimerization was carried out by the addition of staple strands connecting unpaired scaffold domains of the base platform. Gel electrophoresis revealed the dimerization reaction led to one major population (**Fig. S12**), and TEM imaging of these 2×2 L-shaped rotor arrays confirmed the formation of precisely designed objects with four randomly oriented rotor elements (**Fig. 5c, Figs. S13 and S14**). Lastly, introduction of additional DNA strands that bind to rotor arms leaving overhangs capable base-pairing across rotor arms led to formation of the ordered square loops of 4 rotors (**Fig. S15 and S16**). In some cases, formation of individual dsDNA domains between the rotary elements can be visualized, further indicating the interaction between the overhangs (**Fig. S17**). These results establish that interacting arrays of DNA rotors can be feasibly constructed, serving as evidence that rotor arrays exhibiting real order-disorder transitions may be realizable.

**Figure 5:**
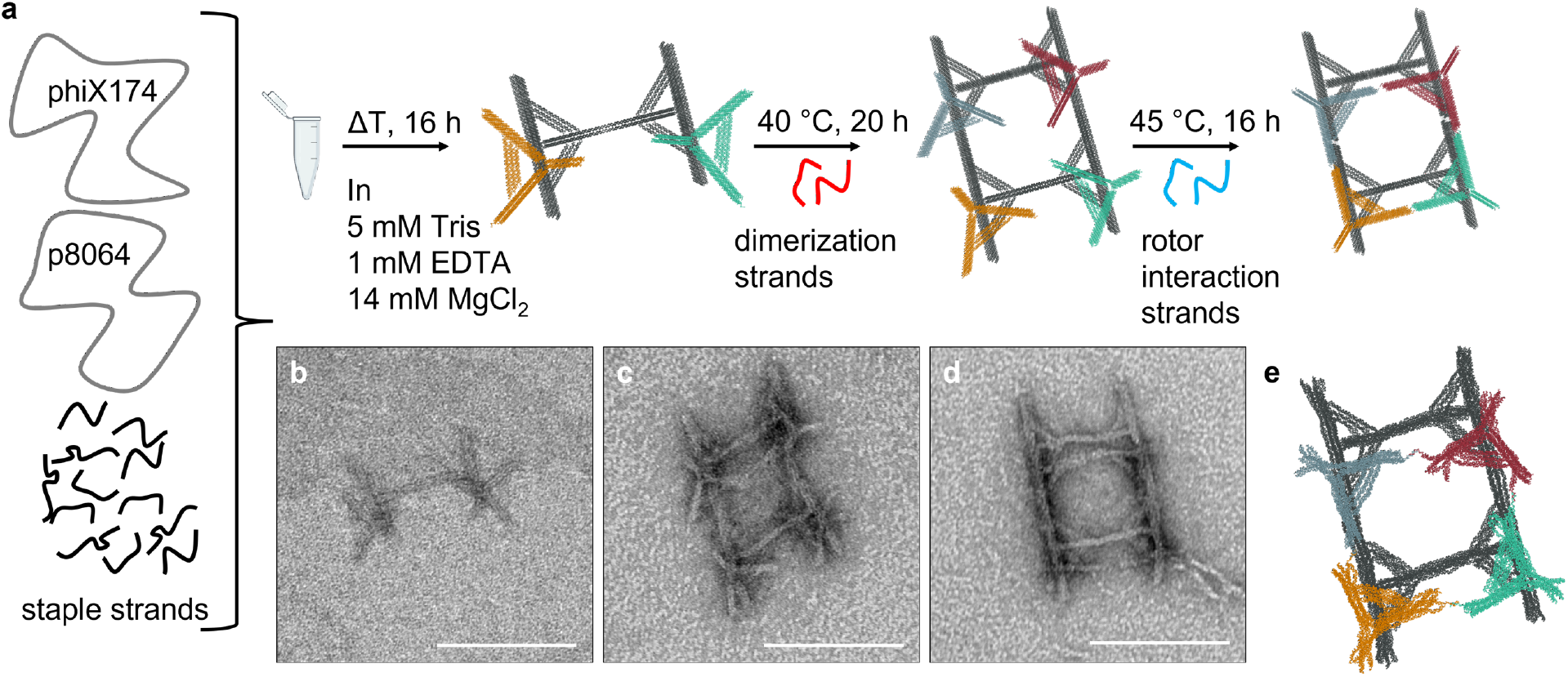
Design, simulation, and fabrication of a small rotor array. (a) Schematics depicting fabrication of double-rotor monomers whose dimerization leads to 2×2 rotor arrays. TEM imaging of: (b) single monomer, (c) 2×2 array displaying a disordered morphology, and (d) 2×2 array displaying an ordered morphology. (e) oxDNA molecular dynamics simulations of an ordered 2×2 array. Scale bar: 100 nm.

## 3. DISCUSSION

This work introduces a new lattice model composed of fixed rotary units that can adopt a discrete set of orientational states and mediate directional interactions with neighboring units. We show *via* MC simulations how these features cause such lattices to exhibit unique order-disorder transition behavior (low transition temperatures and multi-transitions) and captivating ordered morphologies (closed loops, radiating loops, and ladder-like patterns). Although it does not appear that these devices explicitly exhibit conventional second order phase transitions, similarly to the XY model, these systems may require more complicated analysis for the potential existence of phenomena analogous to the vortex transitions described by Kosterlitz and Thouless.^24^ Regardless, a vast design space of rotors remains to be considered. It is possible that some systems, especially those on different lattices such as a Kagome lattice or 3D cubic lattice, could exhibit even more complex and interesting behavior. In addition, while we have studied smaller systems out of experimental interest, system size effects could be significant and warrant further study. Further, it may be worth modifying this model to address a continuum of rotational orientations as opposed to a set of discrete rotational states as we have done here. Finally, implementing multiple types of rotors together in an alternating 2D pattern is experimentally viable; alternating rotor patterns could provide additional interesting morphological behavior.

Crucially, these rotor lattices can be experimentally realized using DNA origami, as we demonstrate for the case of the L-shaped rotors. While we focused on proof-of-concept for feasibility, this experimental work sets the stage for creating larger scale and more diverse sets of rotor lattices whose spatiotemporal ordering behavior can be measured using both modern imaging and fluorescence techniques. The versatility of individual rotor designs could be enhanced using toehold mediated strand displacement^31^ to add and remove overhangs *in situ*. This would effectively modify the rotor interaction tensor of an already-fabricated rotor array; the morphology of existing arrays can potentially be readily switched between the interesting phases that we observed in this study and perhaps more. It is worth noting that the experimental implementation of this system does not correspond perfectly to the presented model, especially when considering neighbor interactions mediated by sticky ends. Attempting to perfectly map the model to the experimental system would involve considering a continuum of states and assigning a temperature-dependent hybridization free energy change^32^ in place of temperature-independent enthalpy terms used in our model.

The rotor lattices introduced here could find future applications in sensing, where local changes in salt or other environmental conditions may be detected on surfaces by local regions of order in large-scale DNA rotor systems using fluorescence or plasmonics. These ordering systems could also be useful for templating inorganic materials in the observed patterns, where such inorganic molecules could be functionalized onto the rotors and subsequently mineralized into robust fixed inorganic surfaces. Rotor lattices could also find use in a pedagogical setting, where basic and accessible imaging like TEM could be used to showcase order-disorder phase transitions to students in a theoretically justified system. Finally, rotor lattices could hypothetically function in a role similar to that of quantum computing^33^: just as quantum computing may be especially useful for solving problem classes whose mathematics are fundamentally similar to the physical behavior of the qubits used to solve them (e.g., *ab initio* electronic structure calculations), these rotor lattices could potentially be used to simulate phase transitions or other mathematical problems involving emergent phenomena from nearest-neighbor interactions. Furthermore, electrical actuation of large-scale DNA rotor systems can lead towards applications like cargo transport.^34^

## 4. CONCLUSION

We developed a model for DNA-based nanoscale rotors which exhibit short-range interactions that can be used to induce large-scale ordering, and which can be switched between ordered and disordered states by modulation of the interaction strength or temperature. This study has elucidated several of the features and achievable order-disorder transitions of this class of DNA rotor devices, where we showed that they are capable of forming several unique morphologies, and individual rotor designs can form up to three different patterns each by way of order-disorder transitions. Finally, we constructed a physical realization of these rotors and demonstrated their capability to assume ordered and disordered morphologies.

## AUTHOR CONTRIBUTIONS

GA and MD conceived the project with help from CEC and MP. MD developed the model, conducted simulations, performed analysis, and prepared the manuscript. WGP performed all experimental work and contributed to the manuscript. BR performed exploratory simulations. CMH designed the experimentally implemented rotors. All authors contributed to manuscript writing and revision.

## ACKNOWLEDGMENTS

This work is supported by the National Science Foundation (Grant No. CMMI-1921955). MD is supported by the National Science Foundation Graduate Research Fellowship (Grant No. DGE-2139754). We acknowledge resources from the Campus Microscopy and Imaging Facility (CMIF) at The Ohio State University for negative stain TEM imaging. AFM imaging was supported by NIH 1S10OD025096-01A1. We acknowledge resources from the Duke Computing Cluster for carrying out oxDNA and MC simulations.

## METHODS

### Simulation of DNA origami rotor arrays

MC simulations were performed on custom scripts written in C++. A single move type was used (perturbation of a single spin). Total system energy was recorded periodically, and specific heat was calculated using Python scripts which can be found in the **data package** for this publication. All simulation codes can be found on GitHub.^35^

### Umbrella sampling

To harmonically constrain this system to an order parameter *χ*, we modify the Hamiltonian to the following:

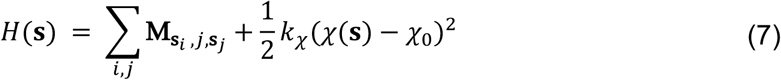

Where *k*_*χ*_ is a harmonic spring constant, generally set to 10 kcal/mol unless this was found to be insufficient; *χ*(**s**) is the instantaneous value of *χ* calculated based on system microstate **s**, and *χ*_0_ is a desired value of *χ* corresponding to the specific umbrella window. Simulations were performed in the range of *χ*_0_ = 0.03 to *χ*_0_ = 1 in increments of 0.03. Free energy was calculated using the Grossfield group’s weighted histogram analysis method (WHAM) software.^36^

### oxDNA simulations

The oxDNA^37^ model and its associated simulation package were used to sample the equilibrium configurational space^38^ of rotor structures and to identify undesired rotation of the rotors. Relaxation of structures was conducted with a constraint on maximum backbone force and with mutual traps between all paired nucleotides to relax the structure into a configuration that could be simulated without these constraints. Simulations were performed at a salt concentration of 0.5 M, equivalent to standard DNA origami fabrication conditions.

### Design and folding of DNA origami nanostructures

DNA origami nanostructures were designed in an iterative process using MagicDNA^29^ and oxDNA^37^. Staple strand routings were fine-tuned in caDNAno. The individual bundles of the DON rotor device are based on a twist-corrected square lattice.^39^ The main bundles of the base-platform are designed to be shape-complementary, allowing seamless head-to-tail multimerization. The struts between the main bundles and the cross-connection help to maintain the parallel orientation of the main bundles. Rotors are connected to the base by two stretches of 4 bases unpaired scaffold DNA, to allow full rotation.^34^ Rotors are designed to carry 0, 1 or 2 overhangs, allowing to tune the strength of interactions. The rotor size allows hybridization of overhangs at the rotors’ interface. Out of plane rotation of the rotor is reduced by the bundle at the rotors back side.^40^

DNA origami nanostructures were folded by mixing all DNA components in folding buffer (5 mM Tris, 1 mM EDTA, pH 8), supplemented with 14 mM MgCl_2_. Scaffold strands (p8064, produced as previously described^41^ and phiX174, purchased from NEB) were used at 10 nM, while all staple strands were used at 60 nM. Thermal annealing was performed by gradually decreasing the temperature from 65 to 20 °C over 16 hours (details in **Table S1**). All staple strands were ordered as desalted products from IDT in IDTE buffer.

Dimerization reactions were performed by pooling purified structures, passivated on the left and right side, respectively, and adding the staple strands designed to link structures head-to-tail. Dimerization strands were added in 10x excess over the DNA origami concentration. Dimerization was performed by incubating the samples at 40 °C for 20 hours. Successful dimerization was confirmed by agarose gel electrophoresis.

Overhangs allowing rotor-rotor interaction were added in a final step. The samples were then subjected to an incubation from 45 – 20 °C over 14 hours. Samples were then imaged without additional purification.

### Atomic force microscopy (AFM)

Gel purified samples were applied onto freshly cleaved mica (V1 mica, Ted Pella) and incubated for 3 minutes. The mica was subsequently rinsed carefully with ddH_2_O and dried using a gentle flow of air. Samples were imaged using a Bruker BioScope Resolve equipped with a Nanoscope V controller using the ScanAsyst in Air mode. SiN probes with triangular cantilevers, a nominal tip radius of 2 nm and a nominal spring constant of 0.4 N/m were used. Scan rates during imaging were usually around 1 Hz.

### Transmission electron microscopy (TEM)

Purified structures were stained using a 1% Uranyl acetate solution and imaged on a FEI Tecnai G2 Spirit electron microscope. Briefly, 6-8 μl of sample were applied on a freshly glow-discharged grid (Electron Microscopy Sciences, Hatfield, PA) and incubated for 5-10 minutes. The sample solution was then removed carefully with Whatman #4 filter paper and the grid immediately stained with two 6 μl drops of 1% Uranyl acetate solution. Grids were dried at least for 20 minutes before imaging.

### Agarose gel electrophoresis

Agarose gel electrophoresis was performed to confirm proper folding of DNA origami nanostructures and to remove excess staple strands and unspecific aggregates. 30 μl of folded DNA origami (10 nM) were run in a 1.5% gel (1.5% agarose, 45 mM Tris, 45 mM boric acid, 1 mM EDTA, 11 mM MgCl_2_) for 120 minutes at 90 V. Gel rigs were submerged into an iced water bath. DNA origami samples were recovered from the gel by excising the target bands, which were subsequently used with Freeze ‘N Squeeze spin columns (BioRad).

