## Supporting Information for "Thermally Reversible Pattern Formation in Arrays of Molecular Rotors"

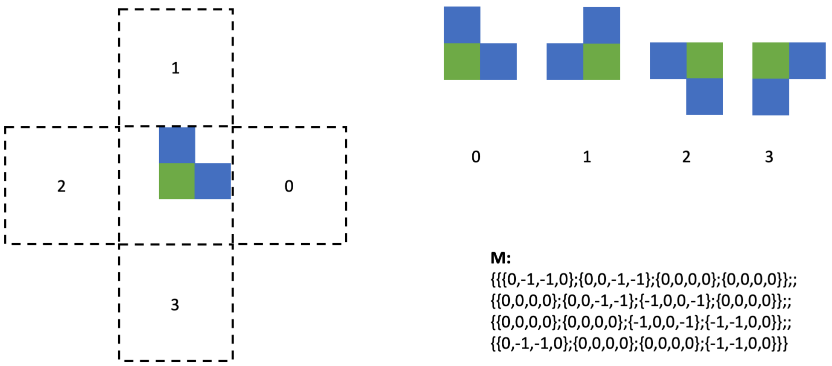


**Figure S1**: Diagram of system states and neighbor indices for L-shaped rotor.

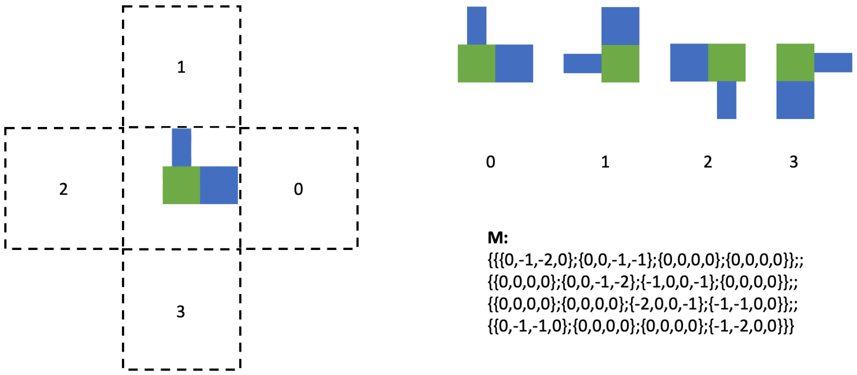


**Figure S2**: Diagram of system states and neighbor indices for chiral rotor.

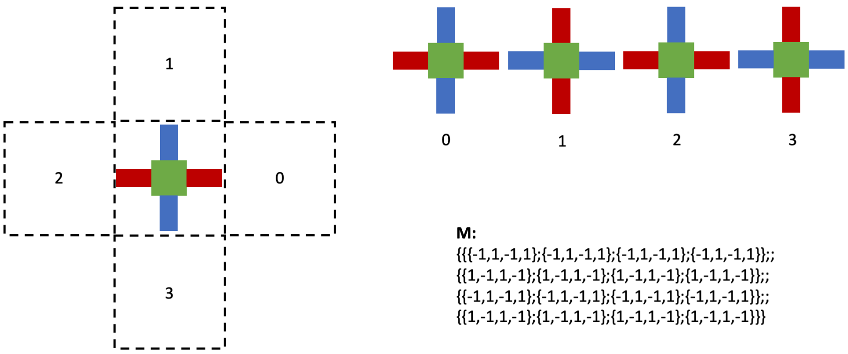


**Figure S3**: Diagram of system states and neighbor indices for 2D Ising-like rotor.

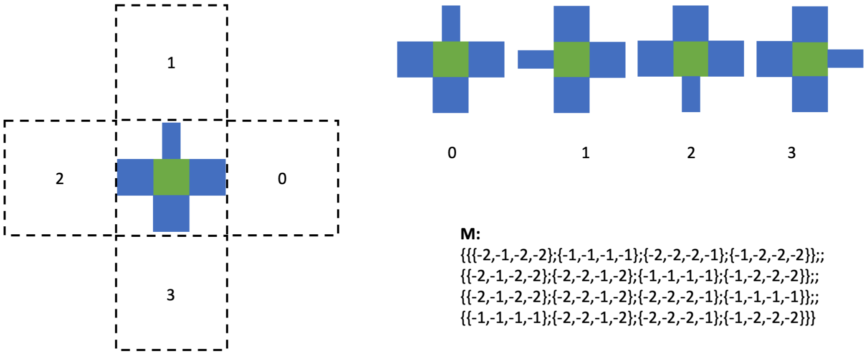


**Figure S4**: Diagram of system states and neighbor indices for 2-2-2-1 rotor.

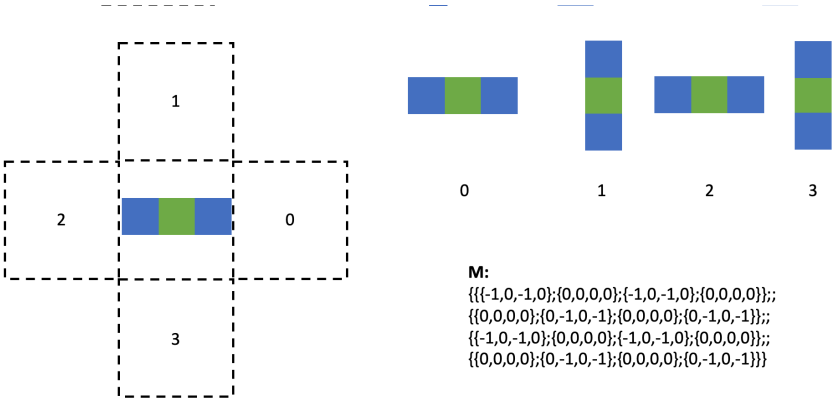


**Figure S5**: Diagram of system states and neighbor indices for quasi-1D Ising rotor.

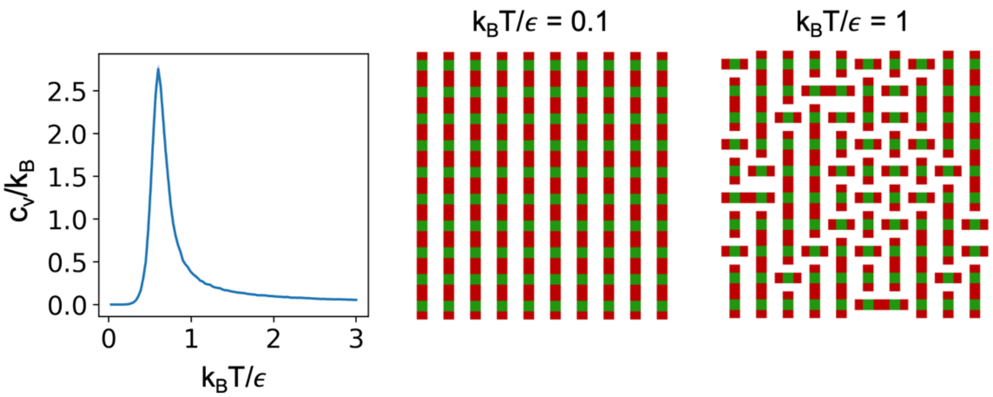


**Figure S6:** Specific heat curve for 10x10 quasi-1D Ising system.

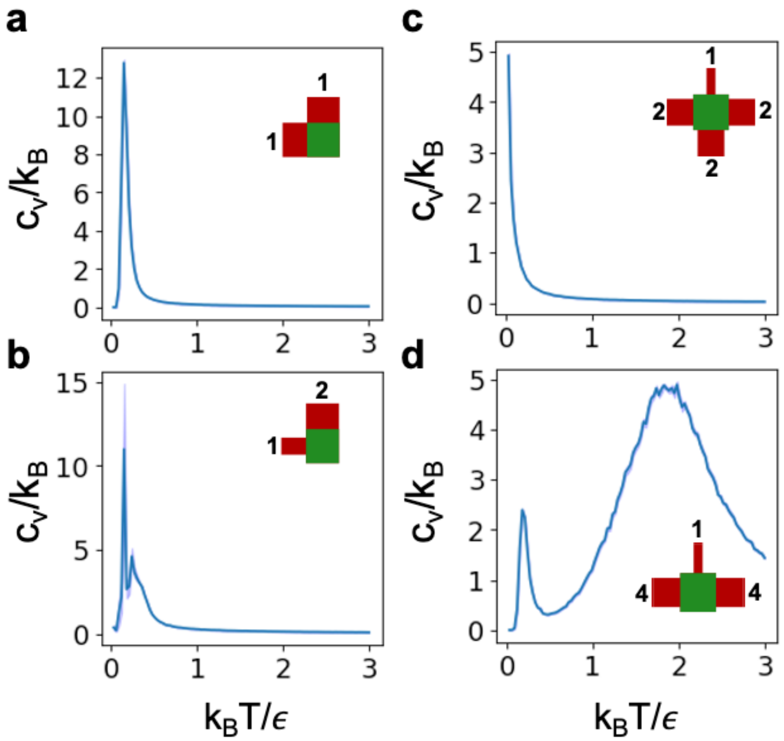


**Figure S7**: Specific heat curves for four designs with no periodic boundary conditions: (a) L-shaped rotor, (b) Chiral rotor, (c) 2-2-2-1 rotor, and (d) 4-1-4 rotor.

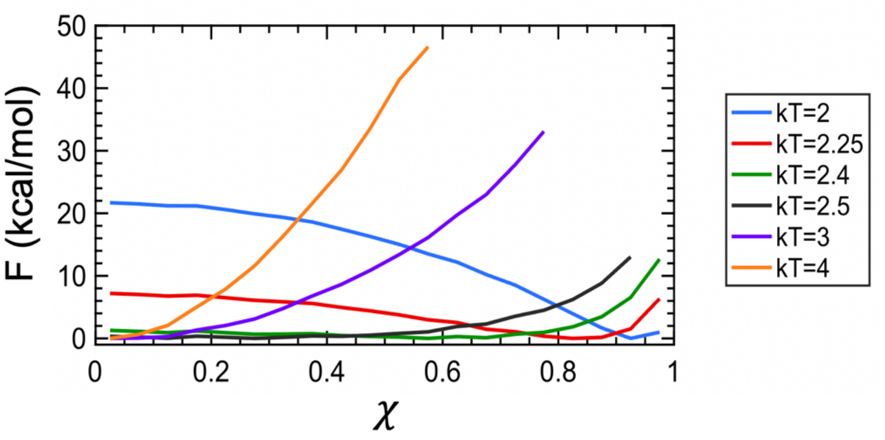


**Figure S8**: Free energy computed as a function of $\chi$ at various temperatures for the 2D Ising lattice using umbrella sampling Monte Carlo simulations.

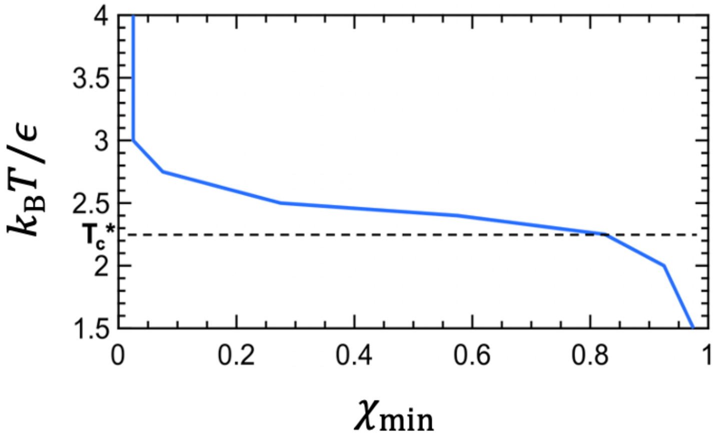


**Figure S9**: Free energy-minimizing value of $\chi$ as a function of temperature, demonstrating a transition in the Ising lattice. $\chi$ acts analogous to the more commonly used order parameter, average magnetization.

**Table S1:** Thermal annealing protocol to fold DON rotor devices.

| **T [ °C]** | **t [min/°C]** |
|---|---|
| 65 | 15 |
| 64-61 | 5 |
| 60 | 8 |
| 59-58 | 12 |
| 57 | 15 |
| 56 | 25 |
| 55 | 36 |
| 54 | 45 |
| 53-49 | 60 |
| 48-45 | 45 |
| 44 | 40 |
| 43-42 | 36 |
| 41-38 | 24 |
| 37-20 | 10 |

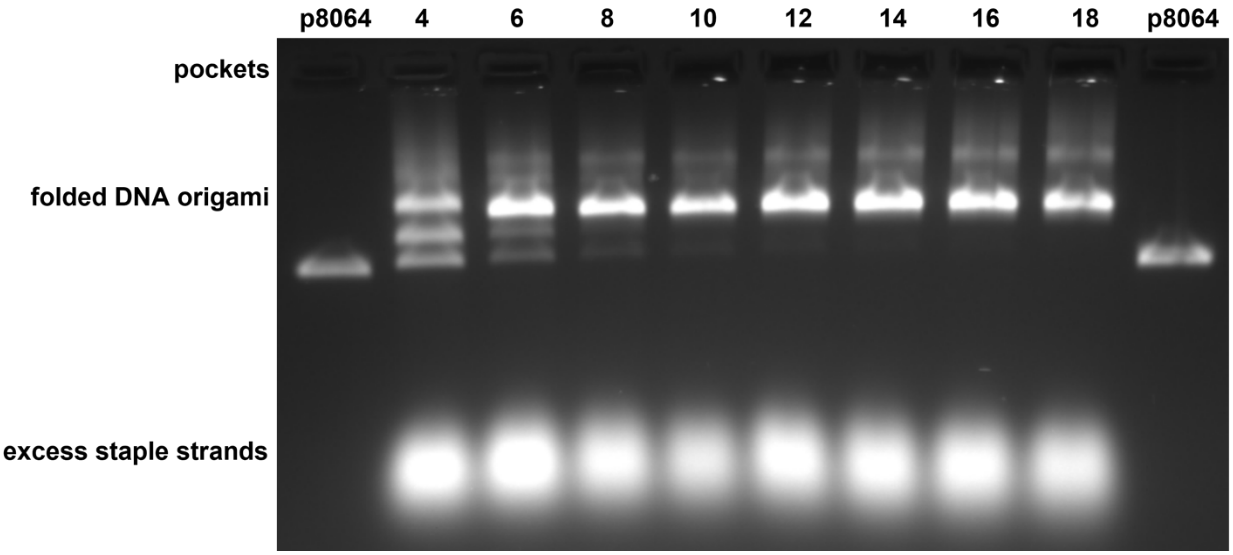


**Figure S10:** Agarose gel electrophoresis of a Magnesium screening, indicating the conditions to properly fold the DON rotor device.

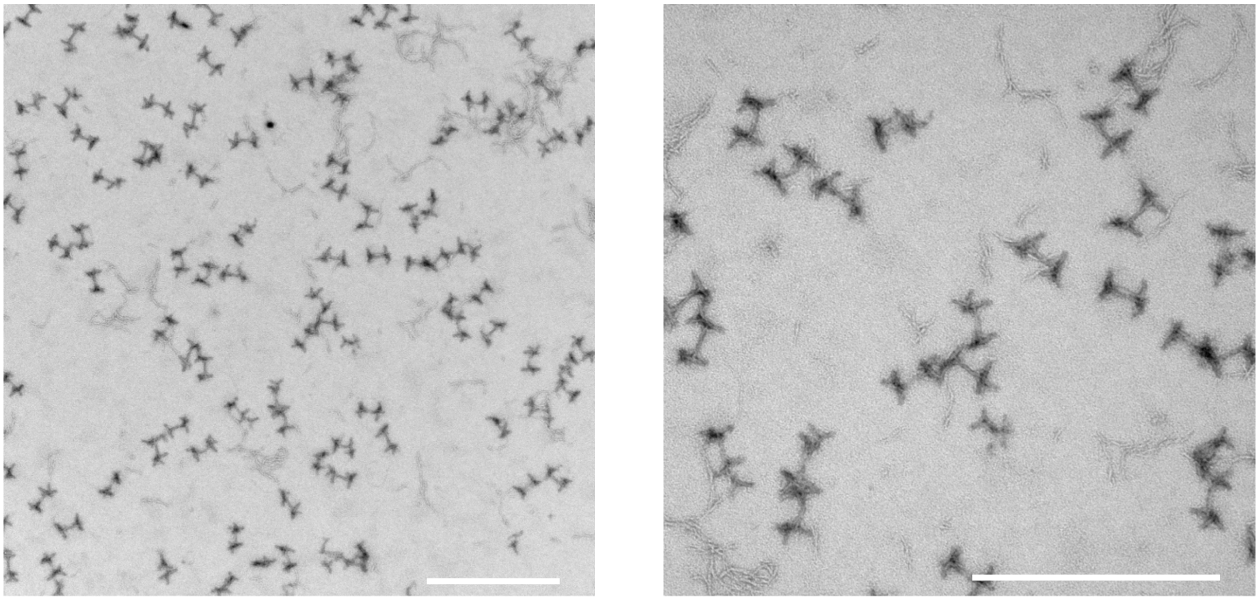

**Figure S11:** TEM micrographs of monomeric DON rotor devices (without overhangs). Scale bar: 500 nm.

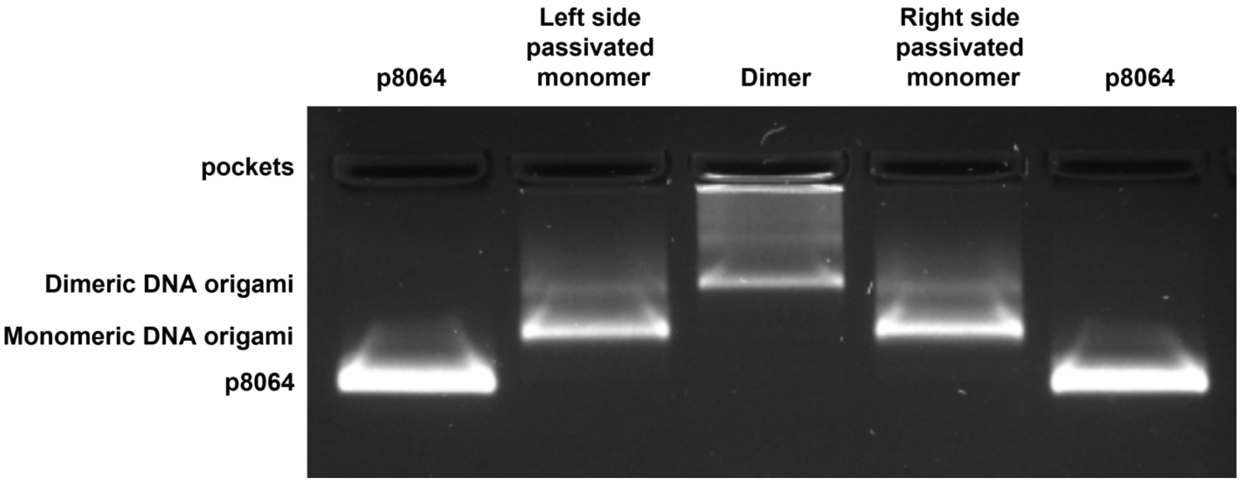


**Figure S12:** Agarose gel electrophoresis of monomers and dimers of the DON. Dimer bands were then excised and extracted using a BioRad Freeze ' N Squeeze kit.

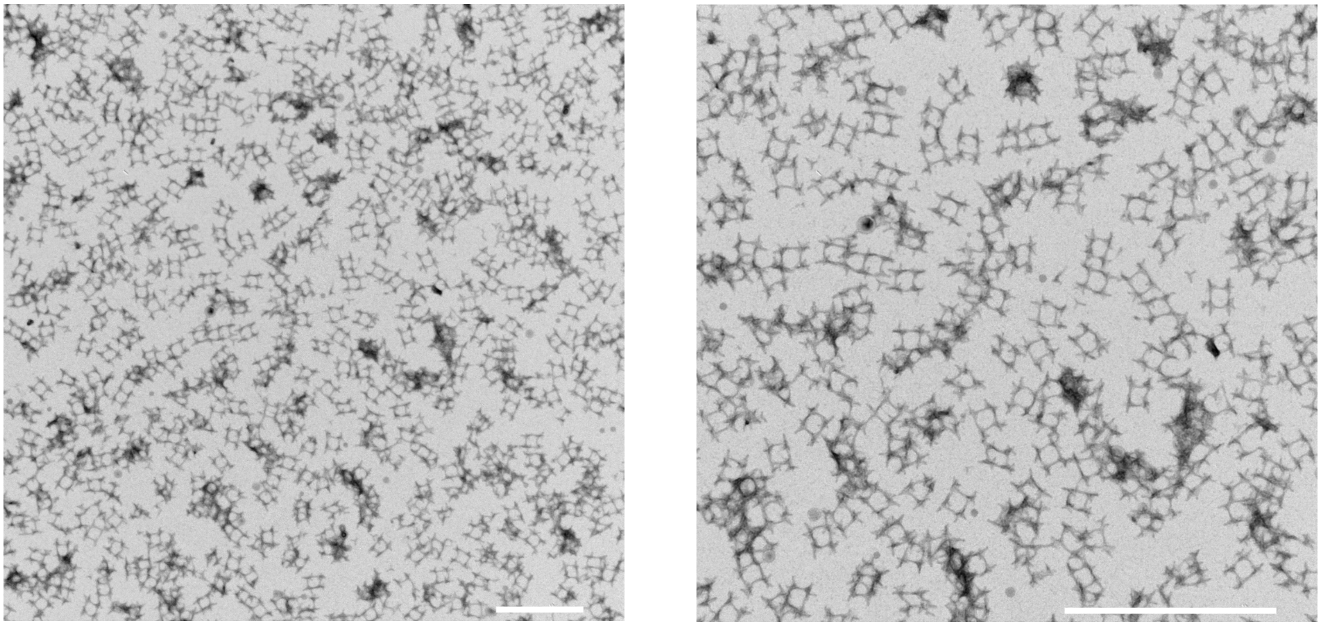

**Figure S13:** Exemplary TEM micrographs of dimeric DON rotor devices. Scale bar: 500 nm.

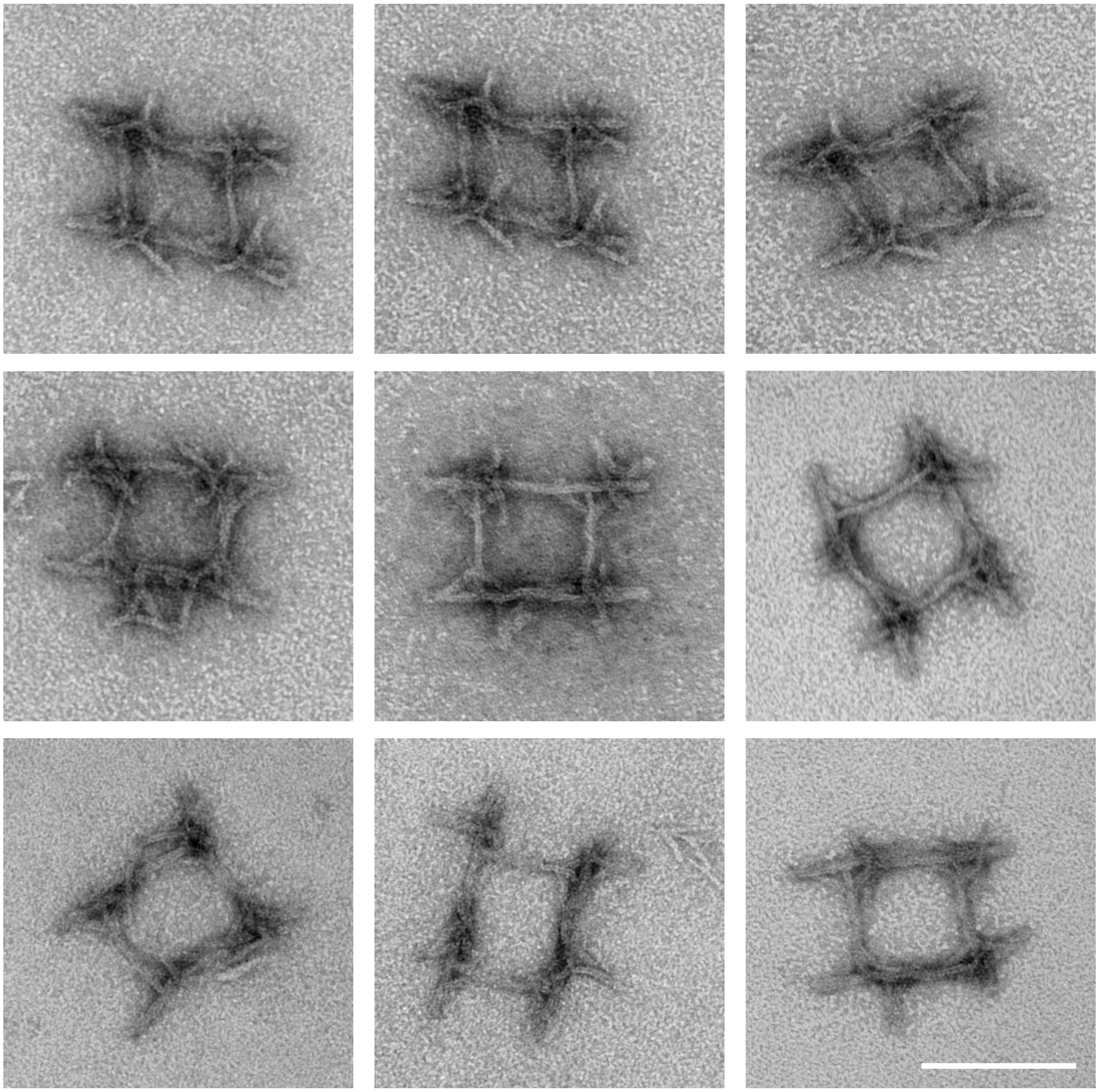

**Fig. S14:** Exemplary TEM images of rotors on 2x2 arrays, showing no alignment. Scale bar: 100 nm.

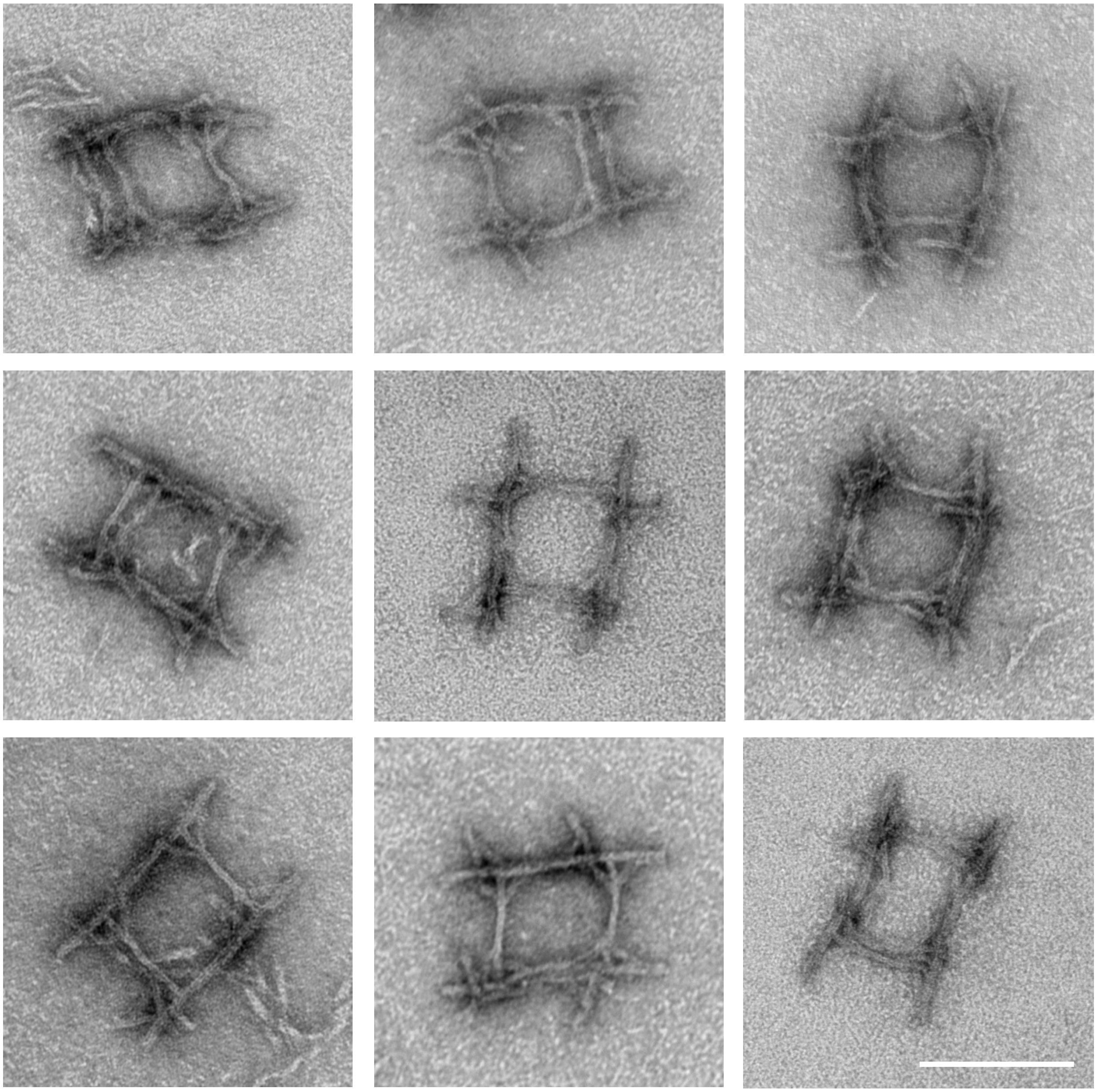

**Fig. S15:** Exemplary TEM images of rotors on 2x2 arrays, showing partial alignment. Scale bar: 100 nm.

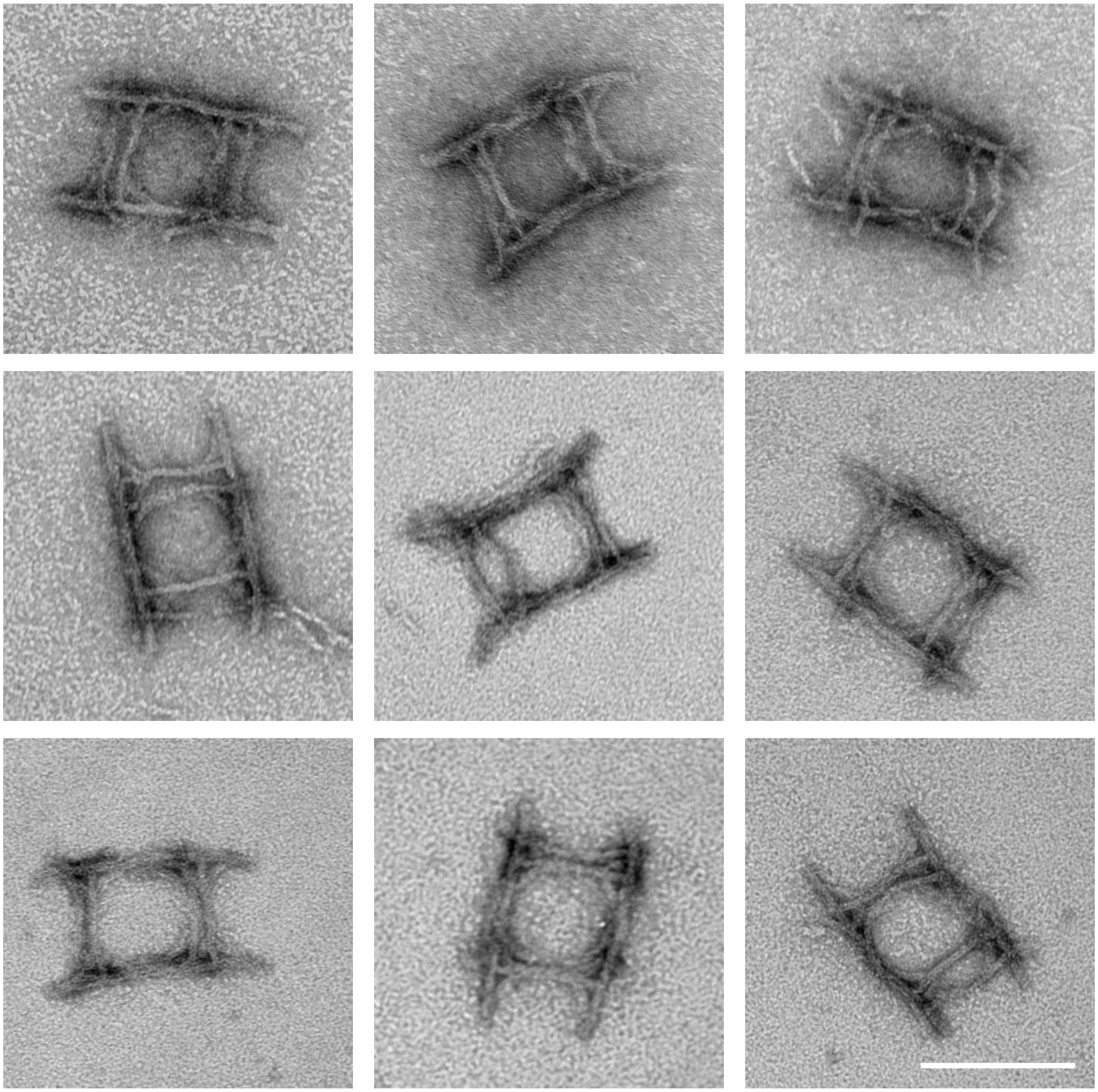

**Fig. S16:** Exemplary TEM images of rotors on 2x2 arrays, showing full alignment. Scale bar: 100 nm.

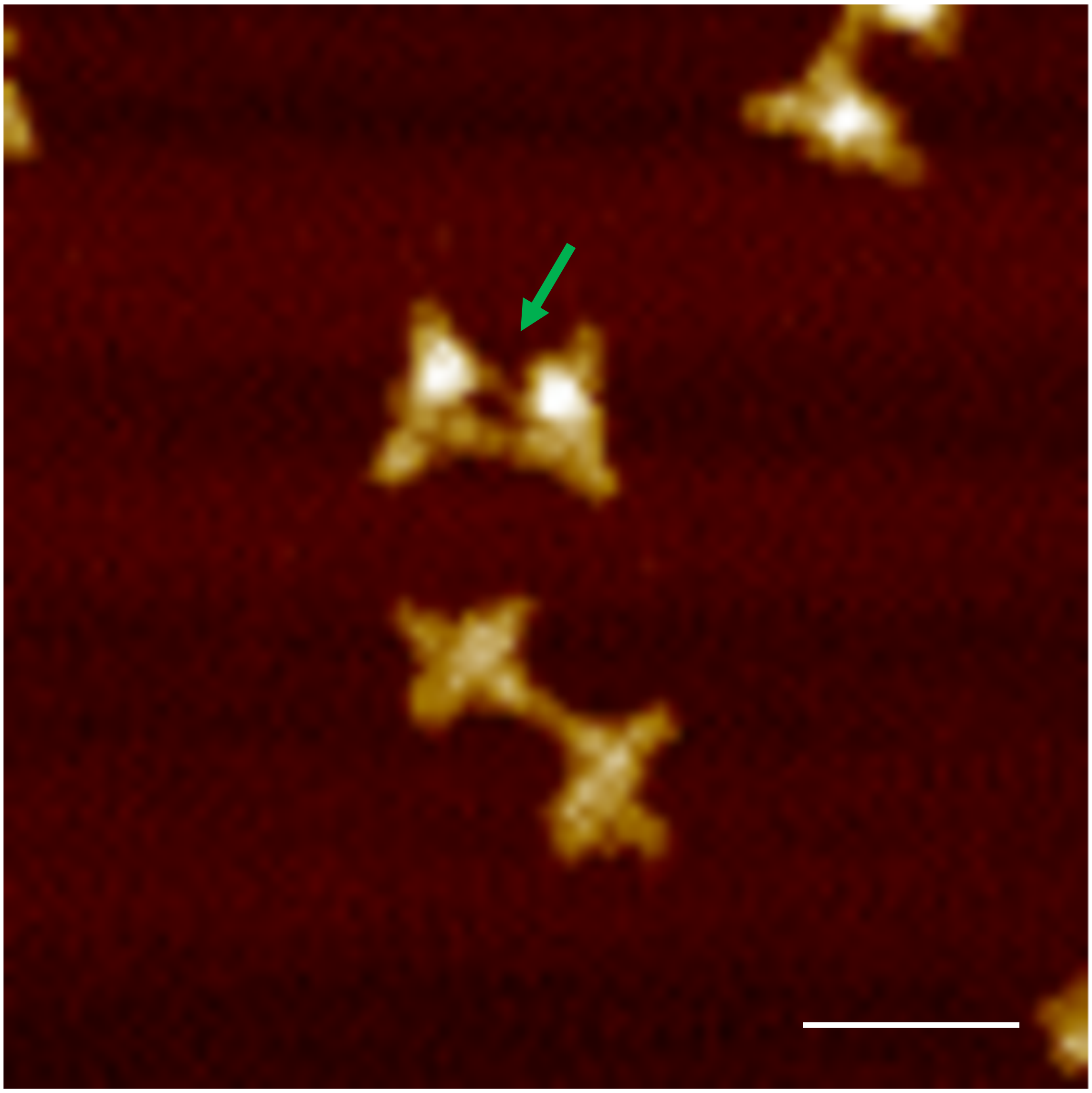

**Fig. S17:** AFM image showing the formed dsDNA connection between the complementary overhangs (green arrow). Scale bar: 100 nm.

**Design:**

The full design of the two-rotor DON will be available on nanobase.org.

**Sequences:**

All oligonucleotide sequences can be found in an Excel spreadsheet in the **data package.**
